# Archaeal, bacterial, and eukaryal microbial community structure of sediment and seawater in a coastal region near Puerto Nuevo, Baja California

**DOI:** 10.1101/324442

**Authors:** Sabah Ul-Hasan, Robert M. Bowers, Andrea Figueroa-Montiel, Alexei F. Licea-Navarro, J. Michael Beman, Tanja Woyke, Clarissa J. Nobile

**Affiliations:** Department of Molecular and Cell Biology, School of Natural Sciences, University of California Merced, Merced, CA, 95343, USA.; Quantitative and Systems Biology Graduate Program, University of California Merced, Merced, CA, 95343, USA.; Department of Energy, Joint Genome Institute, Walnut Creek, CA 94598, USA.; Departamento de Innovación Biomédica, Centro de Investigación Científica y de Educación Científica y de Educación Superior de Ensenada (CICESE), Ensenada, Baja California, México; Department of Life and Environmental Sciences, School of Natural Sciences, University of California Merced, Merced, CA, 95343, USA.

**Keywords:** 16S, 18S, amplicon sequencing, biodiversity, coastal ecosystems, microbial communities, Playas de Rosarito, microbial ecology

## Abstract

Microbial communities control numerous biogeochemical processes critical for ecosystem function and health, particularly in coastal ecosystems. However, comparatively little is known about microbial community structure in coastal regions, such that basic patterns of microbial biodiversity, such as species richness and community composition, are generally understudied. To better understand the global patterns of microbial biodiversity in coastal ecosystems, we characterized sediment and seawater microbial communities for three sites near Puerto Nuevo (Baja California, Mexico) using 16S and 18S rRNA gene amplicon sequencing methods. We found that sediment bacteria, archaea, and eukaryote microbial communities contained approximately 5 × 10^2 fold greater operational taxonomic units (OTUs) than their seawater-based counterparts (p < 0.001). Further, distinct bacterial, archaeal and eukaryal phyla were found in sediment and seawater samples. The phyla Acidobacteria, Chlorobi, and Chloroflexi were found to be abundant and unique to the sediment and Cyanobacteria, Spirochaetae, and Woesearchaeota to the seawater environment. Apicomplexa and Arthropoda were abundant eukaryal phyla found uniquely in the sediment whereas the Cryptomonadales and Protalveolata were detected only in the seawater. Furthermore, bacterial and archaeal communities were statistically different by site (p < 0.05) in both seawater and sediment samples for the Major Outlet site, the site closest to a residential area. In contrast, eukaryal microbial communities were only different among sites in the seawater samples. Overall, these results suggest that our understanding of coastal microbial biodiversity patterns require spatially robust sampling. This study contributes to a growing body of foundational microbial biodiversity and ecology knowledge, providing context to the global change that is induced by urban development.

## Introduction

A surge in marine microbial community studies over the past decade has led to a wealth of new information on the dynamics between microorganisms and their surrounding environments (Fuhrman et al., 2015). As a result, the identification of spatial and temporal patterns of microbial diversity and their correlations to biogeochemical cycling has been vastly expanded (Haskell William Z. et al., 2017; He et al., 2017; Jessup et al., 2004; Kaestli et al., 2017; Kavagutti, 2016; Kirchman, 2016; Prosser et al., 2007; Whitton and Potts, 2007). A recent commentary by Brussaard and colleagues additionally emphasizes the growing role of “big data” from microbial community ecology and biogeochemistry studies in understanding how microbial communities shape the biogeochemical cycling patterns of coasts and oceans (Brussaard et al., 2016). Such information gathered over time provides a path to determining the causes of and the responses to microbial community disturbances (Hunt and Ward, 2015). While these discoveries are innovative in presenting new pieces to the puzzles of marine microbial ecosystems, much coastal microbial diversity is yet to be investigated (Angell et al., 2018; Galloway et al., 2004; Gradoville et al., 2017).

Describing and defining coastal microbial biodiversity, ecology, and associated biogeochemical cycling patterns over time is essential for understanding the impacts that alterations in biodiversity can have on both the environment and human health (Salazar and Sunagawa, 2017). Human-induced environmental impacts, such as the use of antibiotics, nitrogen fertilizers, and other pollutants, can profoundly affect marine microbial biodiversity (Acosta-González et al., 2013; Sun et al., 2013; Wright, 2010). Coastal ecosystems, in particular, are complex interfaces between highly biodiverse sediments and wave action, where pollutants can collect as a result of coastal development and associated human recreational activity. For example, there is evidence of increased rates of infection by beachgoers and surfers resulting from human-induced factors (Leonard et al., 2017). Acosta-González et al. 2013 has additionally demonstrated the impacts of hydrocarbon contamination from oil spills on the bacterial ecology of coastal sediments, where shifts in the microbial communities toward *Gammaprotebacteria, Deltaproteobacteria*, and *Bacteroides* were observed (Acosta-González et 5/15/2018 4:47:00 PMal., 2013). Interestingly, promoting healthy microbial ecosystems can reduce and even reverse the impacts of long-term pollutants (Sarkar and Webster, 2017; Schwarzenbach et al., 2010), broadening the wildlife conservation narrative and giving reason for continued descriptive studies over time and space (Fuhrman et al., 2015).

The Baja California coastline resides within the Southern California Bight marine ecoregion, encompassing the southern California (US) and northern Mexico coast along with vibrant marine biodiversity (Santora et al., 2017; Spalding et al., 2007). The Southern California Bight is a notably important region for biogeochemical cycling due to the presence of strong upwelling events (Santora et al., 2017), yet the southern reaches of the region are comparatively understudied. There are only a handful of microbial biodiversity “omics” studies focused on coastal Baja California, mostly centered on the hypersaline environments of Guerrero Negro (Huerta-Diaz et al., 2011, 2012; Martini et al., 2002; Omoregie et al., 2004; Orphan et al., 2008; Reimer and Huerta-Diaz, 2011; Valdivieso-Ojeda et al., 2014). Here, we describe a case study of coastal microbial biodiversity through ecosystem sampling and sequencing analyses to begin to identify microbial ecology patterns and processes in this understudied region.

In this study, we characterized the bacterial, archaeal, and eukaryal microbial diversity in sediment and seawater of three sites along a 0.45 km range in Puerto Nuevo in Playas de Rosarito, Baja California (Figure 1). We utilized 16S and 18S rRNA gene amplicon sequencing to determine if (1) coastal microbial community richness differs between seawater and sediment environment types among a 0.45 km range to understand the significance of scale, (2) coastal microbial community composition differs between seawater and sediment environment types among a 0.45 km range as an assessment of coastal microbial ecosystems, and (3) similar patterns are observed between bacterial, archaeal and eukaryal microbial communities to assess systems contributions. These findings provide new perspectives on microbial ecology for coastal Baja California, and contribute to strengthening the growing body of work supporting the use of biogeochemical cycling data to predict ecosystem health and homeostasis.

**Figure 1.**
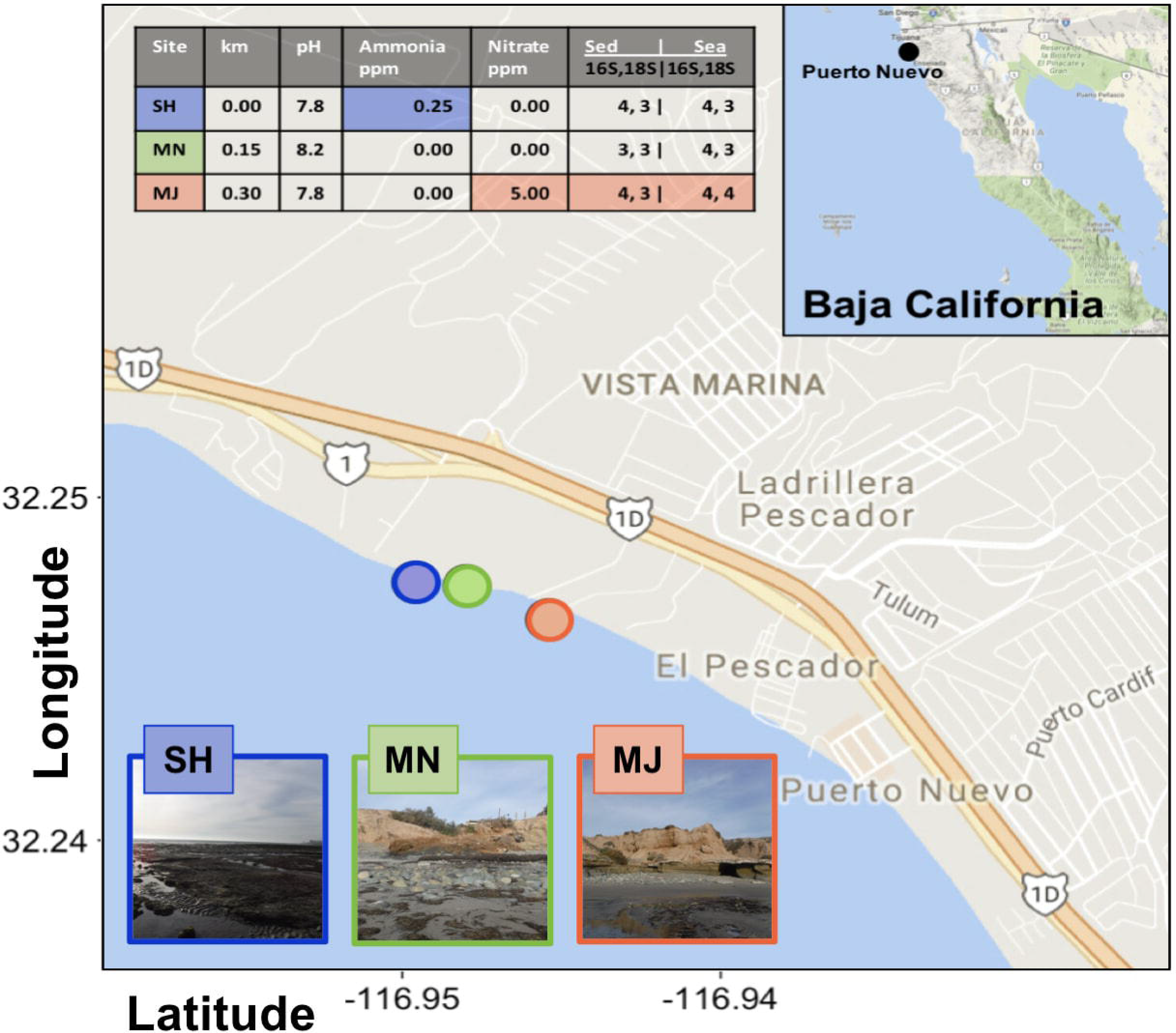
Location and metadata information of sampling sites. Coastal samples were collected at low tide in the summer of 2016 in proximity to Puerto Nuevo, Baja California, Mexico, an avid fishing community near Playas de Rosarito. The three sampling sites are denoted in blue (SH or sheltered), green (MN or Minor Outlet) and orange (MJ or Major Outlet) circles. Four seawater samples and four sediment core samples were collected at each coastal site, for a total of 24 samples for 16S and 18S rRNA gene amplicon sequencing. The Sheltered site is facing a 5-7 m cliff at point 0.0 km, the Minor Outlet site near a small run off outlet or scour at point 0.15 km, and the Major Outlet site is near a large run off outlet and residential area at point 0.3 km. The pH, ammonia (ppm), and nitrate (ppm) levels varied between sites, with differences unique to sites denoted by colored boxes, whereas salinity (1.02 psu) and temperature (~16 °C) were the same. Sequenced samples based on seawater or sediment are displayed in the right-hand table columns for a total of 42 out of 48 possible sequence sets. The inset illustrates the approximate sampling location within Baja California, as denoted with a black circle.

## Materials and Methods

### Sampling

Four biological replicates of 200 mL filtered seawater samples and 8.5 cm length × 1.5 cm diameter sediment core samples were collected during low tide according to previously described methods (Walsh et al., 2015) from three coastal sites near Puerto Nuevo, Mexico between 32.248, −116.948 and 32.246, −116.944 (latitude, longitude) (Figure 1; Supplemental Table 6). Salinity and temperature (°C) were measured for each site along with pH, ammonia (ppm), nitrite (ppm), and nitrate (ppm) using the API Saltwater Master Test Kit. Seawater samples were filtered on-site using sterile 60 mL syringes with 0.1 uM Supor-200 25 mm filters at an approximate rate of 15 mL per min. Filters were then transferred into individual, sterile 2 mL Eppendorf tubes, immediately frozen on dry ice, and stored at −80 °C until further processing. For sediment cores, the tips of sterile 10 cc syringes were cut using sterile razor blades prior to being placed in sediment. Sediment samples were then wrapped with sterile parafilm, immediately frozen on dry ice, and stored at −80 °C until further processing. All samples were handled with sterile nitrile gloves both on- and off-site.

### DNA Extraction, PCR Amplification and Sequencing

DNA from seawater samples was extracted using the QIAGEN DNeasy Blood & Tissue Kit (Qiagen™, Valencia, CA, United States). Filters were cut using sterilized scissors and microbial filter film was homogenized with the Omni Bead Ruptor (Omni International™, Kennesaw, GA, United States) using 0.1, 0.5, and 1.4 um beads. DNA from sediment samples was extracted from 0.5 g of field-moist sediment using the MoBio PowerSoil DNA Isolation Kit (MoBio™, Carlsbad, CA, United States) following the manufacturer’s protocol. Sediment cores were thawed and mixed using sterile weigh boats with sterile spatulas. 0.5 g per sample was used for each extraction. Ribosomal RNA gene amplification was performed for all samples, including a variable 12 bp barcode sequence, following a standard protocol from the Department of Energy Joint Genome Institute (Quast et al., 2013). The V4-V5 region for 16S rRNA of bacteria and archaea (FW 515 F 5’ - GTGYCAGCMGCCGCGGTAA-3’, RV 805R 5’ - CCGYCAATTYMTTTRAGTTT-3’) and the V4 region for the 18S rRNA of eukaryotes (FW 5’ - CCAGCASCYGCGGTAATTCC-3’, RV 5’ - ACTTTCGTTCTTGATYRA-3’) was targeted, with sample validation amplifications to assess extraction quality (Parada et al., 2016; Stoeck et al., 2010). Sequencing was performed on the Illumina MiSeq platform (Illumina™, San Diego, CA, United States) to generate paired-end reads (Caporaso et al., 2012).

### Data Analyses

16S and 18S rRNA reads were recovered from the 24 samples with median lengths of ~380 bp accessible via the <u>Joint Genome Institute Genome Portal</u>. Raw sequences were de-multiplexed and clustered for Operational Taxonomic Units (OTUs) using the iTagger v1.2 and QIIME2 (Bokulich et al., 2017) pipelines for sequence analyses and quality control, with 97% identity or higher via the Silva database (Quast et al., 2013; Tremblay et al., 2015). Identified sequences were then subsampled based on the sample with the lowest read count and parsed according to medium (seawater or sediment) and location (SH, MN, MJ; Figure 1). Sequences were additionally filtered according to matches across biological replicates of six compiled samples (Wat_SH, Sed_SH, Wat_MN, Sed_MN, Wat_MJ, Sed_MJ). After removing contaminant mitochondrial DNA sequences, samples with less than 1,000 paired reads were excluded from analysis. All remaining 16S and 18S rRNA gene sequences were then rarefied at 1,000 OTU reads using the QIIME2 pipeline (Bokulich et al., 2017). Rare phyla were classified by having 10 reads or less of rarefied 1,000.

All statistical tests and visualizations were conducted in R (R Development Core Team 2008) and available via https://github.com/sul-hasan/ (Supplemental Tables 1-5). Changes in microbial community structure were analyzed using non-parametric multivariate analysis of variance (PERMANOVA; Anderson, 2001) using Bray-Curtis distances and Bonferroni p-value correction. Beta diversity differences in community structure were visualized using principal components analysis (PCA) along two axes (Figure 4).

For all univariate data, we used analysis of variance (ANOVA) to determine significant differences among sites, mediums, and site*medium interactions. We used q-q plots and scale-location plots to inspect normality and homoscedasticity, respectively. Where significant differences were detected, Tukey’s Test of Honest Significant Differences was used to determine the range of differences among the sites and interactions.

Boxplots display statistically significant variations in taxa richness (Figure 2), stacked bar charts compare alpha diversity of abundant phyla (Figure 3), PCAs and Venn diagrams compare beta diversity for statistically significant microbial communities (Figure 4), and heatmaps highlight driving taxa by comparison of classes within abundant phyla (Figure 5).

**Figure 2.**
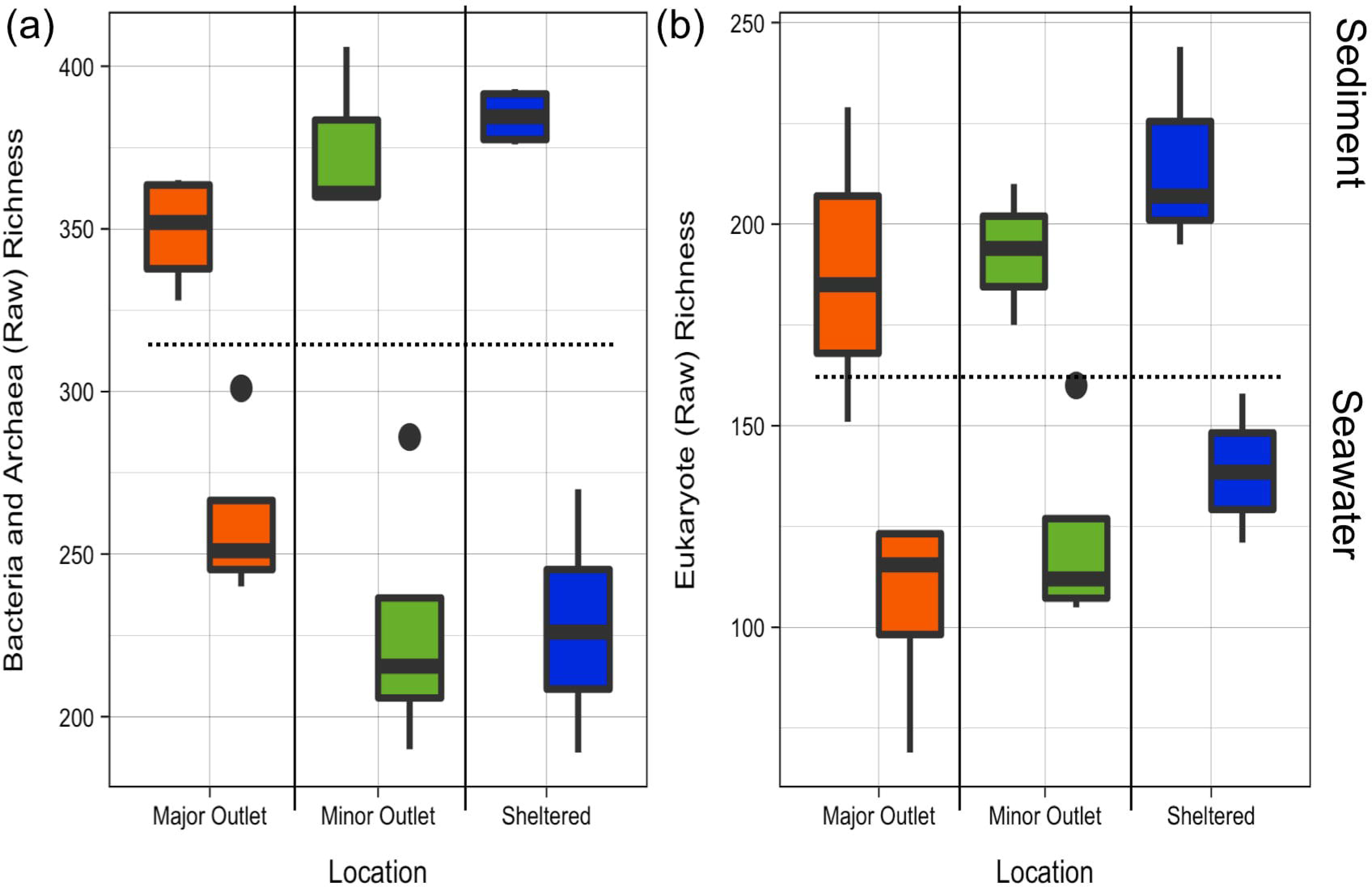
Microbial community richness of sediment and seawater among sites. (a) Boxplot comparisons of rarefied bacterial and archaeal (16S) richness estimates for different environment types (sediment and seawater) and by sampling site (major outlet, minor outlet, sheltered), with a p value (p < 0.001) for environment type. The sediment demonstrates 5.7 × 10^2 fold greater taxa richness relative to seawater. (b) Boxplot comparisons of rarefied eukaryal (18S) richness estimates for different environment types (sediment and seawater) and by sampling site (major outlet, minor outlet, sheltered), with a p value (p < 0.001) for environment type. The sediment demonstrates 4.9 × 10^2 fold greater taxa richness relative to seawater.

**Figure 3.**
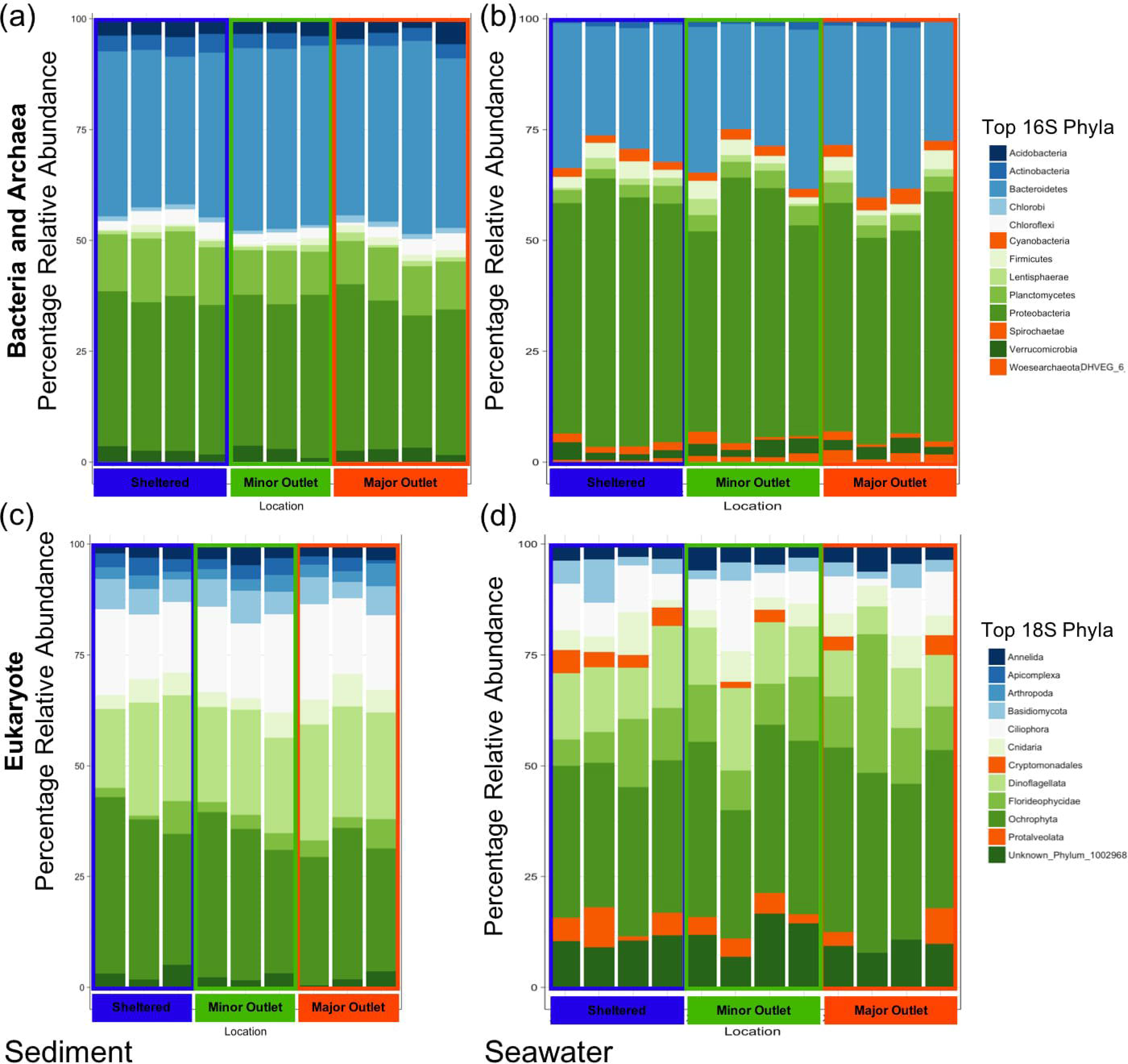
Alpha-diversity of most abundant phylum-level microbial taxa. Biological replicates per sampling site categorized by microbial taxa vs environment type. (a) The top ten bacterial and archaeal phyla in the sediment in order of highest relative abundance to lowest are Bacteroidetes, Proteobacteria, Planctomycetes, Acidobacteria, Actinobacteria, Chloroflexi, Verrucomicrobia, Lentisphaerae, Firmicutes, and Chlorobi. (b) The top ten bacterial and archaeal phyla in the seawater in order of highest relative abundance to lowest are Proteobacteria, Bacteroidetes, Planctomycetes, Firmicutes, Verrucomicrobia, Cyanobacteria, Lentisphaerae, Actinobacteria, Spirochaetae, and Woesearchaeota (DHVEG6). (c) The top ten eukaryal phyla in the sediment in order of highest relative abundance to lowest are Ochrophyta, Dinoflagellata, Ciliophora, Basidiomycota, Cnidaria, Florideophycidae, Annelida, Arthropoda, Apicomplexa, and Unknown Phylum 1002968. (d) The top ten eukaryal phyla in the seawater in order of highest relative abundance to lowest are Ochrophyta, Dinoflagellata, Florideophycidae, Unknown Phylum 1002968, Ciliophora, Cnidaria, Protalveolata, Annelida, Basidiomycota, and Cryptomonadales. Abundant phyla unique to the seawater are highlighted in orange and driving taxa of each significant finding for beta-diversity can be viewed by comparison of OTU tables.

**Figure 4.**
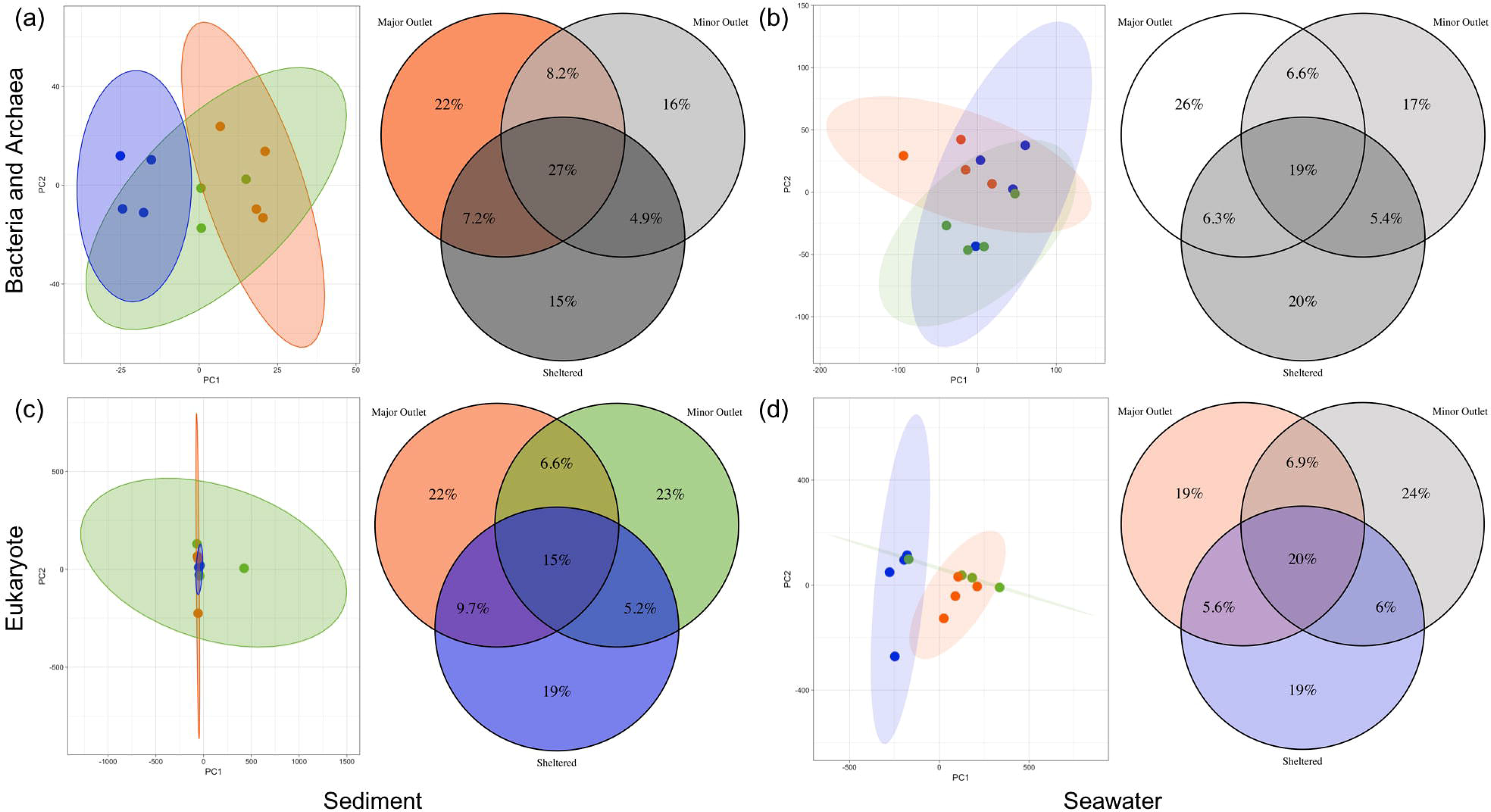
Microbial community composition and beta-diversity. Principal component analyses of PC1xPC2 paired with Venn diagrams of percentage overlapping OTUs for bacterial and archaeal (a-b) and eukaryal (c-d) microbial communities with blue as the Sheltered site, green as the Minor Outlet site, and orange as the Major Outlet site. (a) For the bacterial and archaeal sediment samples, the major outlet site demonstrates a marginal difference when pairwise p-values are compared (p = 0.059 to 0.060), with a 39% PC1xPC2 proportion of variance. (b) For bacterial and archaeal seawater samples, the major outlet is significantly different than the sheltered site (p = 0.025), with a 73% PC1xPC2 proportion of variance. (c) For eukaryal sediment samples, all site locations are significantly different from each other with a 58% PC1xPC2 proportion of variance. (d) For eukaryal seawater samples, none of the site locations differ from each other, with a 79% PC1xPC2 proportion of variance. Shaded ellipses for PCAs encompass 95% confidence. Venn diagrams are colored to indicate significant differences or are shaded in variants of gray to indicate no significant differences.

**Figure 5.**
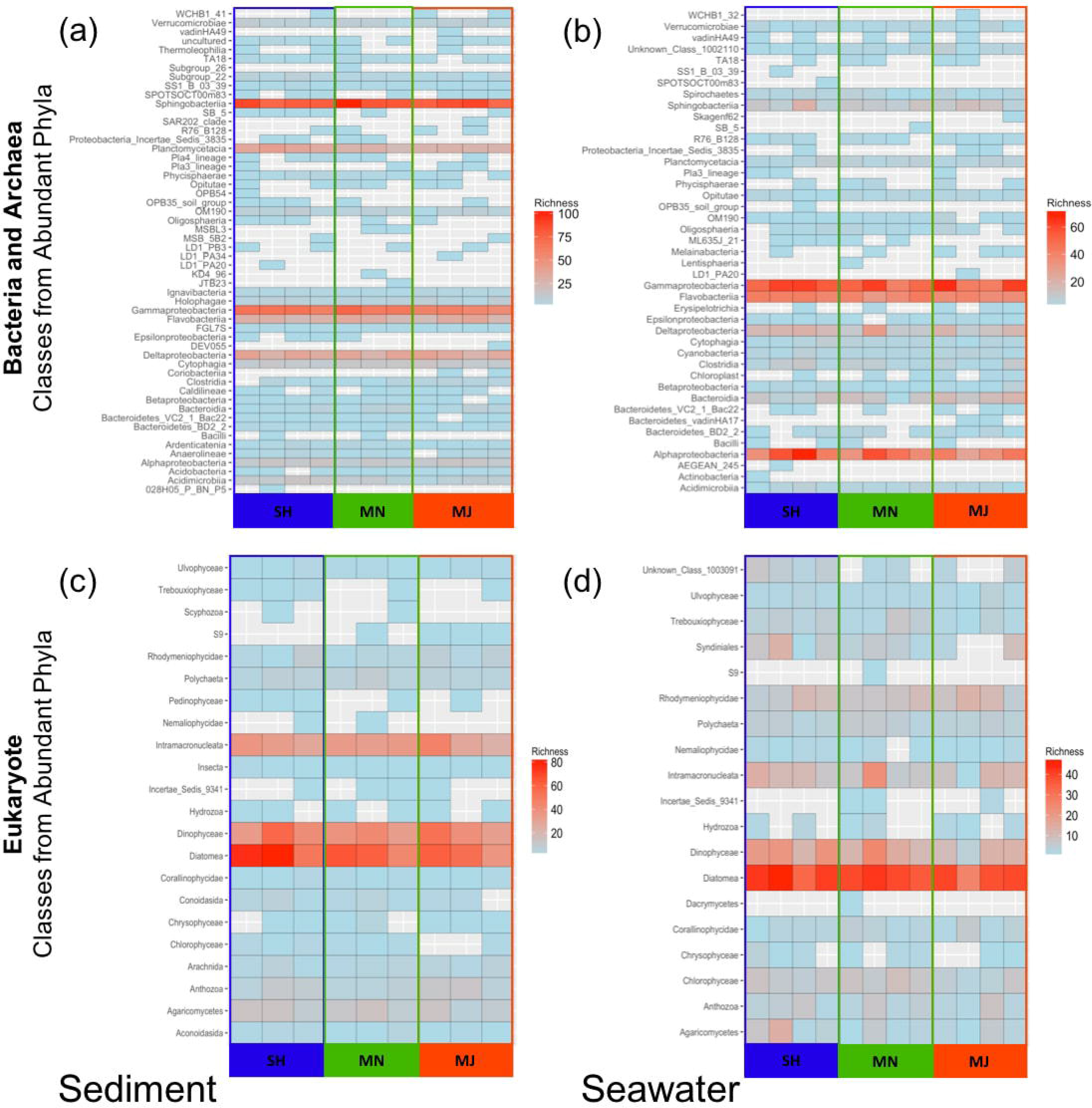
Class richness from abundant phyla. Heat maps of bacterial and archaeal (a-b) and eukaryal (c-d) taxonomic classes from most abundant phyla with blue as the Sheltered site, green as the Minor Outlet site, and orange as the Major Outlet site. Richness is indicated from light blue to red, with gray being a richness of 0, and varies by microbial community category and environment type. (a) The top ten bacterial and archaeal classes from the abundant phyla in the sediment in order of highest relative abundance to lowest are Sphingobacteriia, Gammaproteobacteria, Deltaproteobacteria, Planctomycetacia, Flavobacteriia, Alphaproteobacteria, Cytophagia, Acidimicrobiia, Verrucomicrobiae, and OM190. (b) The top ten bacterial and archaeal classes from the abundant phyla in the seawater in order of highest relative abundance to lowest are Gammaproteobacteria, Alphaproteobacteria, Flavobacteriia, Deltaproteobacteria, Bacteroidia, Sphingobacteriia, Clostridia, Planctomycetacia, Cyanobacteria, and Cytophagia. (c) The top ten eukaryal classes from the abundant phyla in the sediment in order of highest relative abundance to lowest are Diatomea, Dinophyceae, Intramacronucleata, Agaricomycetes, Anthozoa, Polychaeta, Rhodymeniophycidae, Arachnida, Unknown Class 1003091, Conoidasida. (d) The top ten eukaryal classes from the abundant phyla in the seawater in order of highest relative abundance to lowest are Diatomea, Dinophyceae, Intramacronucleata, Rhodymeniophycidae, Chlorophyceae, Syndiniales, Polychaeta, Agaricomycetes, Anthozoa, and Trebouxiophyceae in the seawater.

## Results

### Coastal Puerto Nuevo site locations

Puerto Nuevo is a fishing town located in the municipality of Rosarito, Baja California, Mexico. We sampled three sites within a 0.45 km range off the Puerto Nuevo coastline. The most North-facing site, referred to as the Sheltered (SH) site, contains a 3 m cliff at point 0.0 km and had 0.25 ppm ammonia and 0 ppm nitrate; the site referred to as the Minor Outlet (MN) site near a small run off outlet (or scour) at point 0.15 km had 0 ppm for both ammonia and nitrate; and the site referred to as the Major Outlet (MJ) site near a large run off outlet and residential area at point 0.3 km had 0 ppm ammonia and 5 ppm nitrate. All additional measurements were similar to one another across sampling sites (Figure 1).

### Coastal Puerto Nuevo microbial community richness

Sequencing amplicons for the archaeal and bacterial (16S rRNA gene) and eukaryal (18S rRNA gene) communities resulted in 4,117,060 and 10,019,966 reads, respectively. Reads were then normalized and filtered or rarefied at 1,000 per sample. Sediment archaeal and bacterial communities were found to be >63% richer in OTUs (p < 0.001) than seawater archaeal and bacterial communities for coastal Puerto Nuevo (Figure 2). Similarly, sediment eukaryal communities were found to be >56% richer in OTUs (p < 0.001) than seawater eukaryal communities. These raw results do not account for the mass of the starting samples containing 0.5 g of sediment and 200 mL of filtered seawater, indicating that all microbial communities for sediment are orders of magnitude richer (approximately 5×10^2 fold) relative to those of seawater when calibrated to the amount of sample. Archaeal, bacterial and eukaryal community richness did not differ among sites (p = 0.900 for archaea and bacteria, and p = 0.110 for eukaryotes).

### Coastal Puerto Nuevo microbial community composition

Microbial communities within sediment and seawater environment types revealed specific taxonomic assemblages associated with water versus sediment samples collected from the same locality (Figures 3-5). Similar to OTU richness, microbial community composition differed by medium (p = 0.001). Figure 3 demonstrates how Acidobacteria, Chlorobi, and Chloroflexi are abundant bacterial and archaeal phyla unique to the sediment environment, whereas Cyanobacteria, Spirochaetae, and Woesearchaeota (DHVEG6) are unique to the seawater environment. Within these phyla, Acidimicrobilia, OM190, and Verrucomicrobiae were all unique classes to the sediment and Clostridia, Cyanobacteria, and Bacteroidia were all unique classes to the seawater. Apicomplexa and Arthropoda are abundant eukaryal phyla solely found in the sediment environment, whereas Cryptomonadales and Protalveolata phyla are unique to the seawater environment. Within these phyla, Arachnida, Unknown Class 1003091, and Conoidasida were all unique classes to the sediment and Chlorophyceae, Syndiniales, and Trebuxiophycea were all unique classes to the seawater (Supplemental Image 1).

Figure 4 demonstrates that seawater and sediment archaeal and bacterial communities significantly differed by site (p = 0.016 and p = 0.007, respectively). In both cases, the MJ site significantly differed from the other two (seawater p = 0.003; sediment p = 0.025). Seawater eukaryal communities significantly differed by site (p = 0.040), with all sites significantly different from each other (p = 0.003). Sediment eukaryal communities did not differ significantly by site (p = 0.099).

Rare bacterial and archaeal phyla for sediment with an abundance of 10 or less after rarefaction of 1,000 are Gracilibacteria, Hydrogenedentes, Cloacimonetes, Atribacteria, Marinimicrobia (SAR406), PAUC34f, SHA109, Fibrobacteres, Omnitrophica, TA06, Woesearchaeota (DHVEG6), Deinococcus (Thermus), Elusimicrobia, and Cyanobacteria. Rare bacterial and archaeal classes for sediment with an abundance of 1 after rarefaction of 1,000 are 028H05PBNP5, DEV055, JTB23, KD496, Latescibacteria Incertae Sedis 11404, LD1PA20, LD1PA34, Methanomicrobia, MSB5B2, OPB54, SAR202, Subgroup 26, Thermoplasmata, Unknown Class 1002391, Unknown Class 1002397, and VadinHA49. Rare bacterial and archaeal phyla for seawater with an abundance of 10 or less after rarefaction of 1,000 are Armatimonadetes, Deferribacteres, Gemmatimonadetes, Latescibacteria, Microgenomates, Nitrospirae, Parcubacteria, TA06, Chloroflexi, Elusimicrobia, Omnitrophica, Chlorobi, Deinococcus (Thermus), TM6, Acidobacteria, Cloacimonetes, and Fibrobacteres. Rare bacterial and archaeal classes for seawater with an abundance of 1 after rarefaction of 1,000 are Actinobacteria, Aegean 245, Anaerolineae, Ardenticatenia, Deferribacteres Incertae Sedis 11258, Gemmatimonadetes, Holophagae, Latescibacteria Incertae Sedis 11404, LD1 PA20, Lentisphaeria, Nitrospira, OPB35, SB5, Skagenf62, SPOTSOCT00m83, SS1B0339, Subgroup22, Unknown Class 1002196, Unknown Class 1002468, Unknown Class 1002476, Unknown Class 1002669, and WCHB1_32. Rare eukaryal phyla for sediment abundance of 10 or less after rarefaction of 1,000 are Tunicata, Echinodermata, Platyhelminthes, Ascomycota, Nematoda, SCM37C52, Protalveolata, SGUH942, Mollusca, and Unknown Phylum 1003810. Rare eukaryal classes for sediment with abundance of 1 after rarefaction of 1,000 are Ascidiacea and Polyplacophora. Rare eukaryal phyla for seawater with an abundance of 10 or less after rarefaction of 1,000 are Ascomycota, Brachiopoda, Nematoda, Platyhelminthes, Prymnesiophyceae, Echinodermata, Unknown Phylum 1003810, SGUH942, Tunicata, and SCM37C52. Rare eukaryal classes for seawater with an abundance of 1 after rarefaction of 1,000 are Dacrymycetes, Isochrysidales, Malacostraca, Prymnesiales, S9, Sordariomycetes, and Unknown Class 1003448.

Figure 5 breaks down classes within abundant phyla as a heatmap to identify driving taxa as inferred from Figures 3-4. Sphingobacteria, Proteobacteria Incertae Sedis 3835, and Intramacronucleata are overall richer in the sediment and Gammaproteobacteria, Flavobacteria, Alphaproteobacteria, and Diatomea are overall richer in the seawater. Deltaproteobacteria and Dinophyceae demonstrate richness in both sediment and seawater environment types.

## Discussion

This study focused on the coastal microbial communities of the previously unrepresented location of Puerto Nuevo, Baja California. Our findings support the ideas that: (i) scale matters in demonstrating that coastal microbial communities are dynamic and distinct among sample sites; (ii) spatial variation in composition of sediment microbial communities is stronger than seawater microbial communities, and (iii) the spatial breadth observed may be weaker in the sediment communities as a result of less fluidity in the medium. Our findings that coastal communities differ among sample sites (SH, MN, MJ) and mediums (sediment, seawater) are corroborated by prior studies in other locations (Hao et al., 2016; Langenheder and Ragnarsson, 2007). Furthermore, our observed differences for bacterial, archaeal, and eukaryal microorganisms between sites within a small 0.45 km range raise additional questions related to microbial ecology and biodiversity.

We found that coastal microbial community richness differs between seawater and sediment environment types, but remains unchanged among a 0.45 km range (Figure 2). The sediment environment type is >63% richer for all microorganisms when compared to seawater, which is consistent with recent literature - although less common in archaeal and eukaryal microorganisms (Cleary et al., 2017; Daly et al., 2014; Walsh et al., 2015). Our findings that sediment displays greater microbial richness than seawater could be due to the fact that, similar to soil (Kuzyakov and Blagodatskaya, 2015), sediment also provides a greater surface area for microorganisms to colonize. Microorganisms have more to “grab” (Aleklett et al., 2018) in the sediment and also have access to a higher proportion of food resources from fluctuating fluid debris (Mincer et al., 2016). The sediment environment type also permits an increased likelihood of microbial mats and biofilms to form, which serve important roles in biogeochemical cycling and maintaining ecological homeostasis (Mincer et al., 2016; Nielsen and Risgaard-Petersen, 2015). These two environment types, however, are not mutually exclusive. The seawater environment type is a necessary contributor for refreshing microbial populations within coastal environments. This raises additional questions regarding microbial composition and taxa preference for one environment type versus another.

We observed that coastal microbial community composition differs between seawater and sediment environment types among a small 0.45 km range (Figures 3-5). Interestingly, a relatively recent field study on grasslands investigated bacterial communities between sites ranging from 10 m to 14.4 km (Hao et al., 2015), and concluded that communities vary independent of geographic distance. Discussions of spatial ecology continue to be pertinent for microbial biodiversity in order to understand ecosystem dynamics, with many studies examining soil microbial communities (Ettema and Wardle, 2002; Green and Bohannan, 2006). Much fewer studies, however, focus on coastal microbial communities and the comparisons between short versus long distances between sample sites. Our study on microbial community differences within a 0.45 km range for coastal Puerto Nuevo highlights the heterogeneity of coastal sediment microbial communities.

We observed distinct patterns between bacterial, archaeal and eukaryal microbial communities (Figure 3-4). Interestingly, we observed vastly different patterns dependent on the environment and the microbial sequence type (Figure 4). Acidobacteria, Chlorobi, and Chloroflexi were found to be abundant bacterial phyla unique to the sediment environment. Acidobacteria is one of the most abundant phyla found on Earth, and is known to be more competitive in soils, which may indicate why it is observed in higher abundances in sediment over seawater (Kielak et al., 2016). Chlorobi and Chloroflexi are photosynthesizing bacterial taxa that demonstrate niche contributions to the sediment, such as sulfur cycling (Camanocha and Dewhirst, 2014, 2014; Wasmund et al., 2016); this contrasts to the preference of photosynthesizing Cyanobacteria in seawater (Gao Yonghui et al., 2014; Makhalanyane et al., 2015; Paerl, 2017). Spirochaetae and Woesearchaeota (DHVEG6) are also abundant bacterial and archaeal phyla unique to the seawater environment, although their ecological roles are largely unknown. Woesearchaeota (DHVEG6) was the only abundant and unique archaeal phyla in each environment type. Apicomplexa and Arthropoda are abundant eukaryal phyla unique to the sediment environment, which is consistent with the fact that Apicomplexa are common marine parasites (Frénal et al., 2017) and that many Arthropoda invertebrates burrow and reproduce in the sediment (Burgess et al., 2015; Nedelec et al., 2014). In contrast, the photosynthesizing Cryptomonadales and Protalveolata eukaryal phyla were found to be unique to the seawater environment. Taken together, our findings inclusive of bacterial, archaeal and eukaryal microorganisms, fill a knowledge gap in our understanding of microbial biodiversity patterns in an understudied region.

Understanding the intricacies of biodiversity is critically important for conservation and human health. The expanding inclusion of microbial biodiversity as part of the larger biodiversity conversation serve as important pieces to Earth’s microbial ecology puzzle (Colwell, 1997; Gilbert et al., 2014; Mishra, 2015). In this investigation, we have expanded our understanding of microbial diversity and community composition in a near-shore marine environment - an environment type that has been generally understudied. Our analysis of coastal microbial communities north of Puerto Nuevo, Baja California, which combined 16S and 18S rRNA gene sequencing approaches of coastal seawater and sediment, identified strong relationships between sampling sites and environment types, consistent with previous studies (Green and Bohannan, 2006; Schimel, 1995). Our findings also highlight the importance of spatial and temporal sample continuity in understanding microbial biodiversity patterns, and provides context to the global change that is induced by urban development.

## Acknowledgements

We thank researchers at the Ensenada Center for Scientific Research and Higher Education (CICESE), UC Merced researchers Dr. Jesse Wilson, Nicholas C. Dove, Dr. Stephen C. Hart, and Michael E. Malloy, and researchers Lisa J. Cohen and Jessica M. Blanton.

## Author Contributions

SU performed laboratory work and statistical analyses and RMB assisted with statistical analyses. SU, AFM, ALN, JMB, and TW contributed to the experimental design. SU, RMB, AFM, ALN, JMB, TW, and CJN contributed to the writing of the manuscript.

